# A mammalian inferior colliculus model for sound source separation using interaural time differences

**DOI:** 10.1101/2024.10.30.621075

**Authors:** Christian Leibold, Sebastian Groß

## Abstract

The inferior colliculus (IC) is a central hub in the ascending auditory brainstem. It hosts many neurons tuned to interaural time differences (ITDs). ITD tuning, however, is already observed and generated one synapse upstream in the superior olivary complex and the physiological mechanisms as well as the functional purpose of the IC projection remain partially unresolved. Here, we argue that combining ITD sensitive inputs from medial superior olive (MSO) and lateral superior olive (LSO) requires a temporally well adjusted delay of cross-hemispheric fibers from LSO to IC, given the fast synaptic kinetics of IC neurons. We present a normative model of the midbrain auditory circuitry that finds an optimal cross-hemispheric delay of 0.3 cycles and optimal synaptic strengths by maximizing the firing rate of IC neurons for a stimulus at a given ITD. The model suggests that, by varying the relative synaptic weight of MSO and LSO input, individual neurons are optimized to transmit information of all sound sources in a complex auditory scene. ITD tuning of IC neurons would then results as a side effect. The model focuses on the low-frequency range, is consistent with the distribution of best ITDs observed in experimental recordings and performs close to optimal in sound source reconstruction.

**Author Summary:** Natural acoustic scenes consist of many spatially distributed and concurrently active sound sources. We show how such concurrent sounds can be optimally reconstructed solely using information about interaural arrival time differences. We argue that this optimal solution is almost matched by a neural circuit model inspired by the ascending auditory pathway. The model is computationally very efficient because changing only one synaptic weight moves the azimuthal focus of a midbrain neuron. The model predicts a universal phase delay of cross-hemispheric delay lines in the ascending auditory brainstem.

## Introduction

Separation of sound sources in noisy acoustic environments is one of the most intriguing capabilities of the human auditory system [1] with artificial sound separation solutions reaching good performance only very recently (e.g. compare [2] and [3]). In search for the underlying biological neural mechanisms of speech processing and/or detection in noise, binaural hearing has been suggested to play an important role [4–10]. Neural mechanisms of binaural processing in the ascending auditory pathway are generally discussed in the context of the nuclei at which binaural interactions take place first [11–13]: The medial and the lateral superior olive (MSO, LSO). The majority of the available data about the neural encoding of the two main binaural cues, interaural time difference (ITD) and interaural level difference (ILD), however, stems from the higher-order inferior colliculus (IC) (e.g. [14–17]). Mostly, binaural tuning in the IC is then interpreted as a proxy of the original representation in MSO and LSO. Although there are notable exceptions that aim to explain discrepancies between IC and superior olivary binaural tuning characteristics in the light of the local circuitry [18–20], and even suggest mechanisms of de-novo generation of ILD sensitivity [21], a normative functional approach of binaural representations in the IC based on circuit mechanisms is missing. Traditionally, the IC has been assumed to be “sluggish” [22] implying that IC activity establishes a rate code requiring long averaging windows and assuming that all fine-scale temporal structure is lost. This assumption has meanwhile been disproved with stimulus fine structure observable up to almost about 1 kHz in electrophysiological recordings [23–25]. These in-vivo findings also match the relatively fast feed-forward excitatory synaptic kinetics in the IC of about 2 ms [26].

In this paper, we present a normative theory of how IC neurons read the brainstem activity of MSO and LSO, assuming that both nuclei represent ITDs as two independent populations, each one of which only conveys the summed activity of all its neurons in a frequency band further downstream [27–33]. We argue that best ITDs of IC neurons can thereby vary over a whole hemisphere with only an adjustment of the relative synaptic weight (LSO vs. MSO) to be required. In contrast to most normative models of sound localization that maximize localization acuity [30, 34, 35], the present theory optimizes sound source separability of mixtures of sounds. It thereby naturally accounts for binaural unmasking, a well-known and long-studied effect showing the binaural improvement of detection thresholds in absence of binaural signal-to-noise differences [36, 37], and for which ITDs are the dominant cues at low frequencies [38].

Binaural unmasking has so far attracted only little attention of biologically inspired modelling [33, 39, 40], despite highly evolved psychophysical models [41–45]; for review of classical models see [46]. The present theory suggests the temporally fine-tuned interactions of afferent ITD representations at the level of the IC play a key role for the separation of low-frequency sounds, combining mechanisms of binaural unmasking and sound localization.

## Models and Methods

An auditory scene is a superposition of sounds arriving from multiple sources. Typically, only one source *S*(*t*) is considered as the signal to be extracted. In this paper, the signal is assumed to have constant ITD Δ*t* (*>* 0 for sounds on the right), and the same sound intensity at the two ears (an approximation for low-frequency sounds). Thus, sound signals arriving at the right *R* and left *L* ear are linear combinations

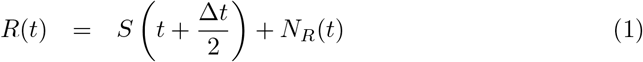

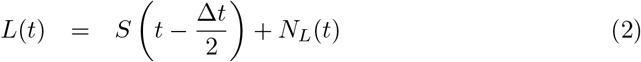

where *N*_*R/L*_ summarize the contributions of the non-signal sounds. Hence, the eardrums translate two pressure waves *R* and *L* into neural activity that encodes three unknowns *S, N*_*R*_ and *N*_*L*_.

Minimizing the mean square error between a linear estimate 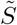 of *S* from the two “measurements” *R* and *L* requires inverting the mixture matrix *A*, which combines equations (1) and (2), and is easiest expressed in the frequency domain, where delays Δ*t/*2 translate to phases e^i*ω*Δ*t/*2^,

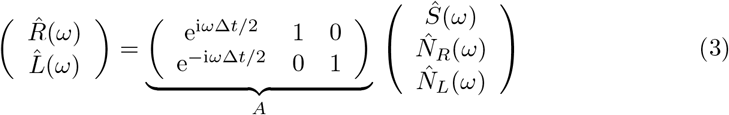

The mean square optimal reconstruction [32] is obtained by the Moore-Penrose pseudo inverse *A*^*^ = lim_*δ*→0_(*A*^†^ *A* + *δ*𝟙)^−1^ *A*^†^ [47], with † denoting the Hermitian conjugate. The frequency representation in the limit *δ* → 0,

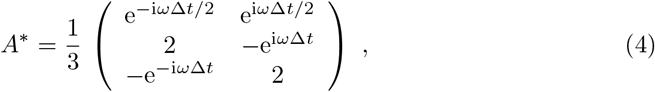

corresponds to the time domain representation of the optimal estimates

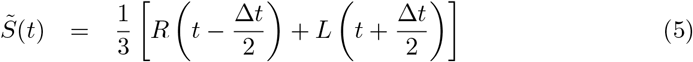

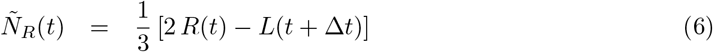

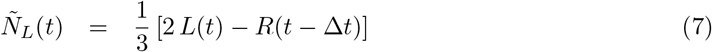

Eq. 5 can be interpreted as Jeffress’ delay-line model for a neuron in which the internal delay between left and right ear exactly compensates the ITD, i.e., a Jeffress-type coincidence detector neuron with the delay offset *d* optimally (in the Euclidean sense) reconstructs sound sources originating from an ITD Δ*t* = *d*.

Delay lines, however, have not been found in the mammalian sound localization circuitry [48], and thus alternative solutions are of interest of how Eq. (5) can be implemented in ways more consistent with mammalian brainstem anatomy.

Sensitivity to ITDs and submillisecond synaptic integration has not only been reported for MSO neurons, but also for LSO neurons [52–54], with pure tone-ITD sensitivity in the low-frequency range [55]. Whereas MSO neurons receive excitatory and inhibitory inputs from both ears, LSO neurons combine ipsilateral excitation with contralateral inhibition. In the simplest form one thus may model the activity of right-hemispheric MSO and left-hemispheric LSO as

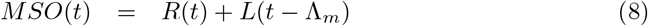

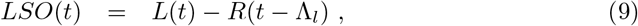

with Λ_*m/l*_ denoting the cross-hemispheric latencies for the projections to MSO/LSO. Notably, *R* and *L*, strictly speaking are no longer pressure waves but neural activity. For the sake of analytical tractability we, however, assume that phase-locked neural inputs approximately match the sound wave if summed over a large set of axonal fibers, thereby interpreting negative values as below spontaneous activity. High numbers of synaptic release sites per axon at a single MSO cell justify this approximation [56].

MSO and LSO activity are combined in the inferior colliculus (IC) where principal excitatory pathways converge from the contralateral LSO and the ipsilateral MSO (Fig. 1 and [57, 58]), i.e., for IC of the right hemisphere one finds,

**Fig 1.**
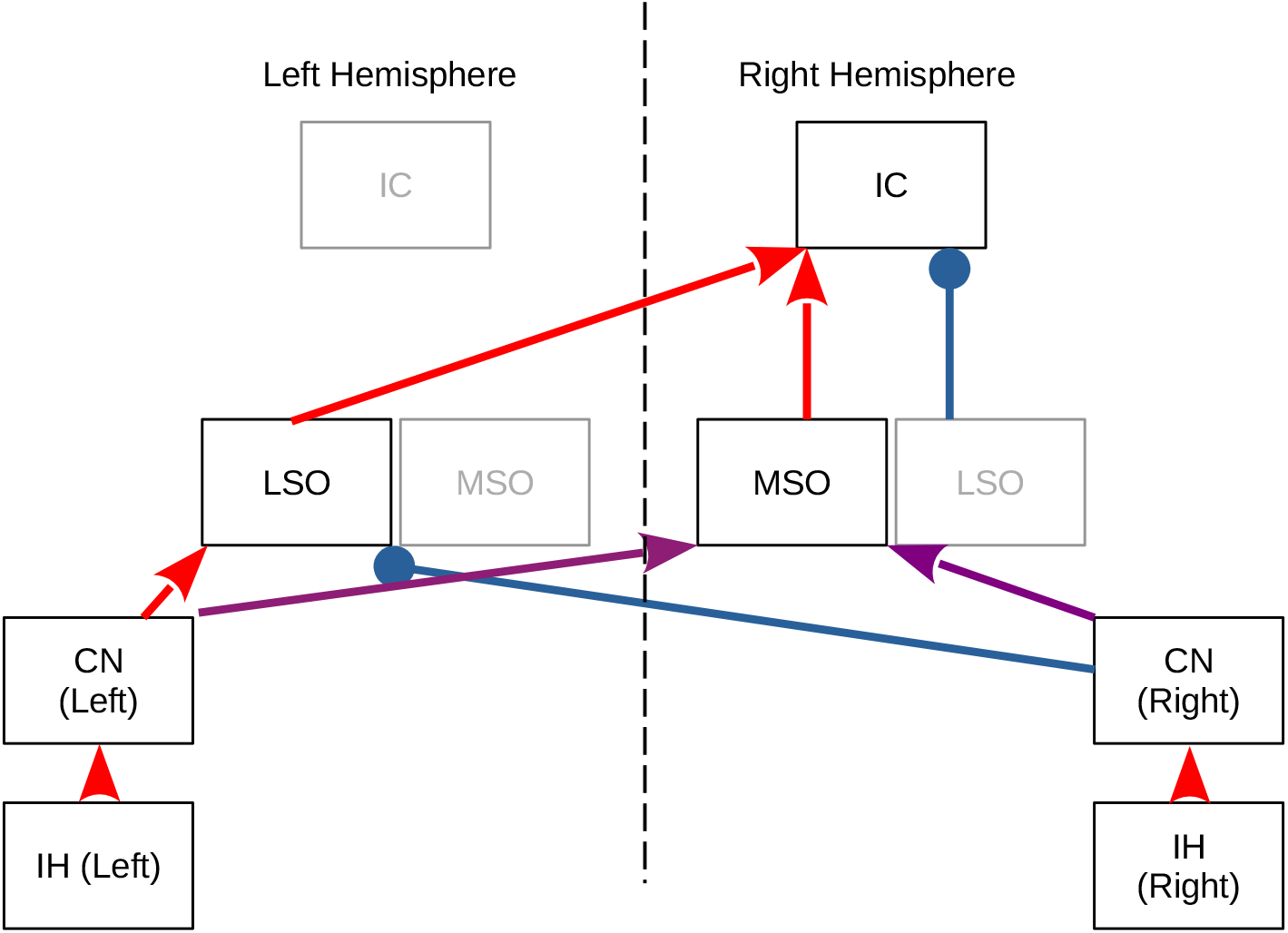
Ascending binaural brainstem anatomy. The inner hair cells (IH) convert mechanical vibrations of the basilar membrane into an electrical signal that is conveyed to the ipsilateral cochlear nuclei (CN) via glutamatergic synaptic transmission (red) of spiral ganglion neurons. CN bushy cell axon innervate bilateral MSO and LSO neurons either directly via glutamatergic synapses (red) or indirectly via glycinergic synapses (blue) from the medial nucleus of the trapezoid body. Whereas for LSO neurons, the presynaptic neurotransmitters separate according to hemisphere, MSO neurons receive mixed (purple) glutamatergic and glycinergic [49–51] synapses from each hemisphere. IC neurons receive glutamatergic synapses from ipsilateral MSO and contralateral LSO, as well as glycinergic synapses from the ipsilateral LSO. The model in this paper is devised for the right IC. Due to symmetry a left IC model can be obtained by flipping right and left ear input.

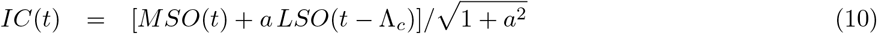

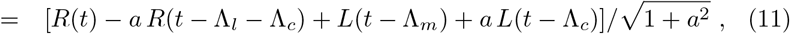

with *a >* 0 denoting the relative synaptic strength and Λ_*c*_ the relative cross-hemispheric (commissural) delay of the contralateral LSO input to IC (via the stria of Held). The factor 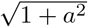 normalizes the sum of squared weights to 1, removing effects on the output firing rate induced by the total synaptic drive. Notably, the present model design allows for binaural processing already in one hemisphere (see also [31]), which is in contrast to standard two channel approaches as e.g. [27, 33, 59], and which would account for the conserved localization ability in the unaffected hemisphere of patients with midbrain stroke [60, 61].

So far our theory neglects frequency-specific processing, and thus, in order to increase biological realism, Eqs. (8-11) should be considered in a best frequency (BF)-channel dependent manner. The Fourier transform of Eq. (11) reads

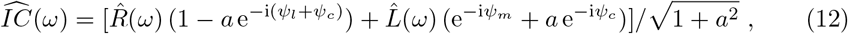

with *ψ*_*m/l/c*_ = 2*πω* Λ_*m/l/c*_ denoting the phase delays introduced by the respective latencies Λ. We next intend to optimize the response of the IC neurons in frequency band *ω* to an isolated sound source and therefore set *N*_*R/L*_ = 0. Accordingly, we set 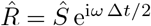 and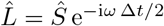, and thus the amplitude of the IC response equals

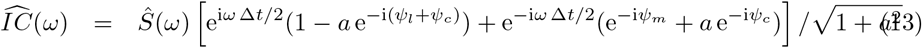

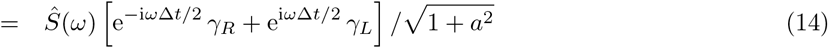

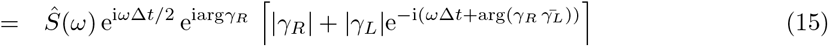

with

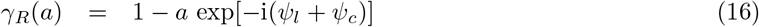

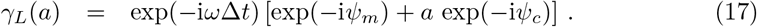

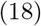

The ITD tuning curve of an IC neuron in the frequency channel *ω* = 2*π* BF is then modeled by the power of the Fourier coefficient,

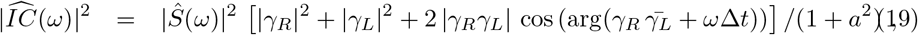

An advantage of the frequency domain representation is that the contralateral transmission delays Λ_*l,m,c*_ can be allowed to vary with frequency *ω* and therefore implement any type of frequency-dependent phase *ψ*(*ω*) = Λ(*ω*) *ω*, including characteristic phases and delays.

In this paper, we will constrain the commissural latencies presynaptic to MSO and LSO to their average experimentally reported values (*ψ*_*m*_ = 0.125 (cyc) [62]; *ψ*_*l*_ = 0 [55, 63]) and maintain *ψ*_*c*_ as a free parameter.

Moreover, we also allow the LSO synaptic weight *a* to take negative values reflecting the ipsilateral inhibitory projection [57], which would change the above formulas to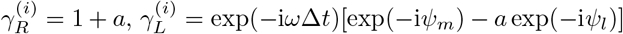, and evidently is independent of the commissural phase delay *ψ*_*c*_.

The dependence of the ITD tuning curves |*IC*| ^2^ (Eq. 19) on the synaptic weight *a* is illustrated in Fig. 2A for two different frequency channels and two commissural phases *ψ*_*c*_. Increasing *a* amplifies the LSO contribution and can effectively shift the best ITD (position of the peak firing rate) to the left or to the right, depending on the relative commissural phase delay *ψ*_*c*_. The shift in best ITD/best IPD via a change in the relative synaptic weight *a* occurs gradually in some intervals of the *a* axis with larger shifts for lower best frequencies, whereas the dependence of best ITD on *ψ*_*c*_ appears more complex (Fig. 2B) but hints at the existence of a low-*ψ*_*c*_ and a high-*ψ*_*c*_ regime. Notably, for *a* = −1, the MSO and LSO contribution driven by the right ear input exactly cancel out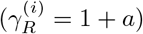, and so no ITD information remains and the tuning curve is flat.

**Fig 2.**
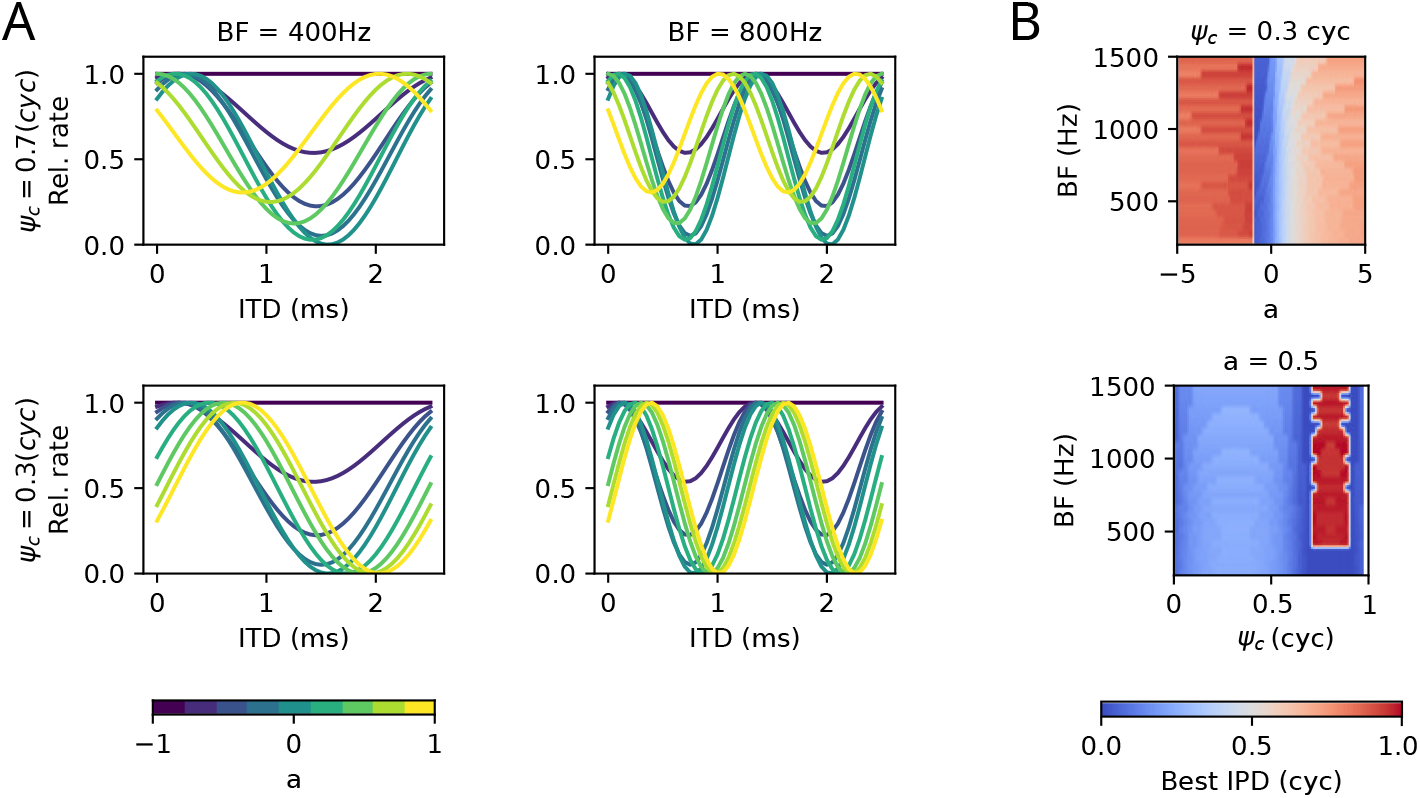
Example ITD tuning curves. (A) ITD tuning curves derived from Eq. (19) are shown for four exemplary conditions and varying relative LSO weights *a* (colors). Top panels are obtained with a large commissural phase *ψ*_*c*_ = 0.7 cyc, bottom panels for small commissural phase *ψ*_*c*_ = 0.3 cyc. Left and right column correspond to best frequency channels of 400 and 800 Hz, respectively. (B) Best interaural phase differences (IPD = ITD × frequency) are color coded for fixed *ψ*_*c*_ = 0.3 cyc as a function of *a* (top) and for fixed *a* = 0.5 as a function of *ψ*_*c*_.

A systematic way to evaluate the influence of the two free parameters *a* and *ψ*_*c*_ is to search for optimal combinations.

Here, a two-step optimization process was performed independently in each BF channel. First, for a given target ITD and *ψ*_*c*_, the optimal weight

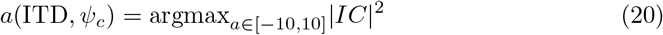

was chosen to maximize the firing rate of an IC cell according to Eq. (19) for stimuli at a target ITD value. This objective deviates from the usual approach to represent auditory space using a wide range of best ITDs, but rather tries to transmit at maximal rate for a signal at the target ITD. In other words, independently of the azimuthal location of a sound source, there should be an IC neuron that is able to convey sound information at maximum rate.

In a second step *ψ*_*c*_, was determined by minimizing the loss function

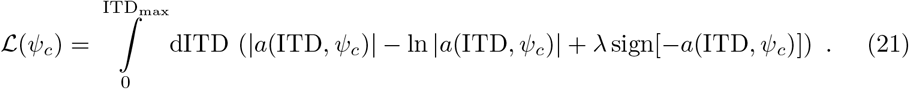

By construction, the loss function tries to minimize the mean relative LSO weight *a*(ITD) (accounting for the relatively small low-frequency LSO), while the − ln term avoids pure MSO solutions with *a* = 0. The regularization *λ* sign[−*a*(ITD, *ψ*_*c*_)] punishes negative weights *a* by means of the sign function, i.e., it punishes solutions including the inhibitory ipsilateral pathway, which, from anatomy, is known to be much smaller in size [57]. Results in this paper were obtained with regularization factor *λ* = 1, the coupling strength *a* that maximizes the objective is further referred to as *a*_*opt*_.

The outcomes of the optimization process are illustrated in Fig. 3 for three different values of maximal ITDs. The largest ITD bound ITD_max_ = 0.7 ms roughly corresponds to human head diameters (and will be used for most further simulations), 0.3 ms approximates the situation in cats, 0.15 ms roughly matches the physiological ITD range in gerbils. The optimal parameters are remarkably similar for all three choices of ITD_max_, with a smooth dependence of the optimal *a* only on the product IPD = ITD × BF. Negative LSO weights *a <* 0 only seem necessary for large head sizes and high BF. Interestingly, the best commissural phase is at around 0.3 cycles under all conditions.

**Fig 3.**
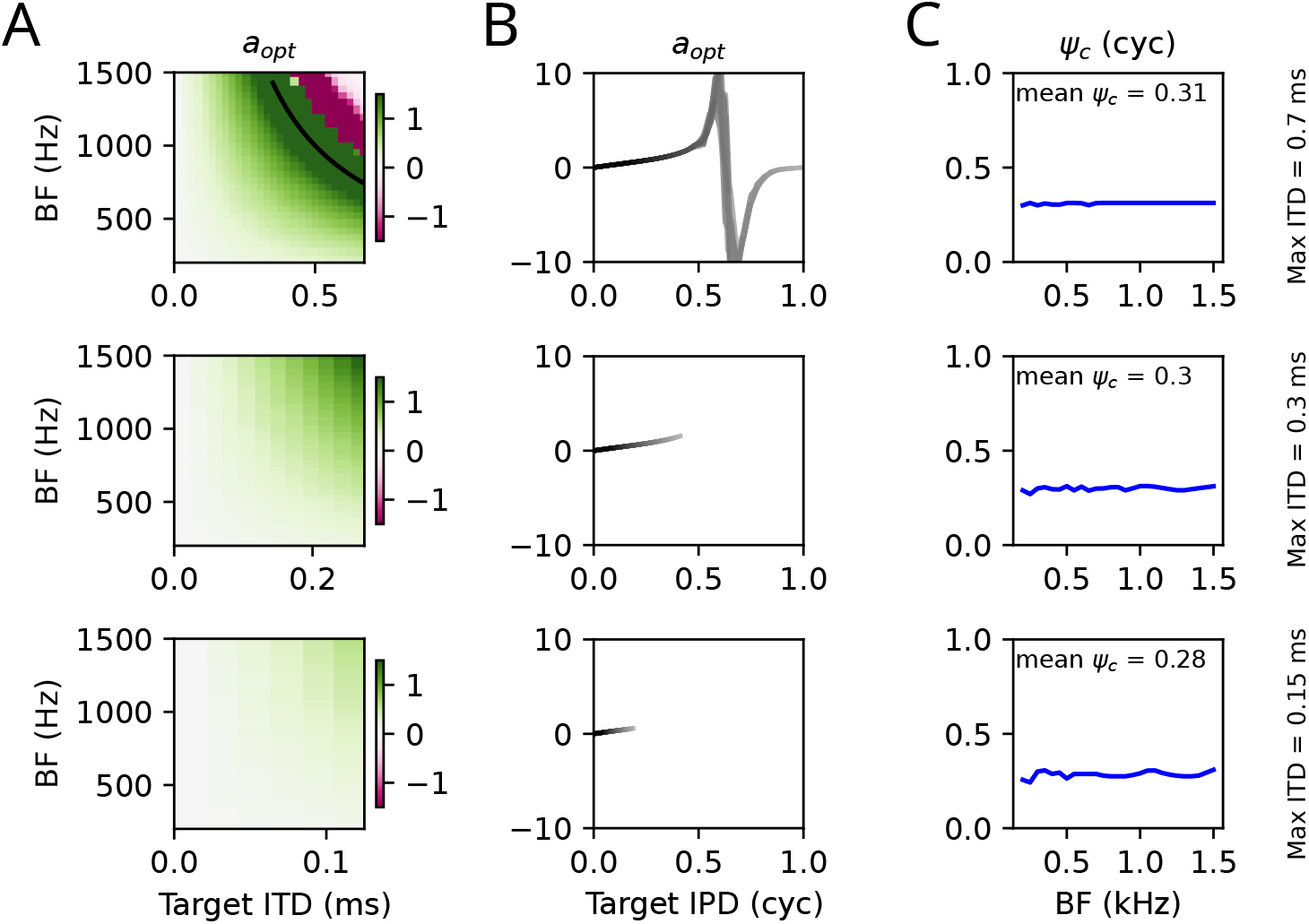
Optimal parameters. The optimal parameters *a* (A: color coded as function of target ITD; color truncated at |*a*| = 1.5; B: as a function of target IPD; grey levels indicate different BF channels from 200 Hz, dark to 1500 Hz, bright) and *ψ*_*c*_ (C) for transmitting a stimulus at a given target ITD/IPD at maximum rate as they were obtained by the optimization procedure described in the main text. Rows show optimal parameters for different maximal physiological ITD (as indicated by x-axis on the left and text on the right). The black line in the top left panel indicates the *π*-limit, i.e., BF×ITD= 0.5.

In order to understand the mechanisms underlying the optimal parameter choice, the input space of the IC (MSO and LSO activity) is examined in Fig. 4 for pure tone stimuli at the BF of three frequency channels and varying ITDs. The trajectories in input space strongly depend on the commissural phase. For the optimal value *ψ*_*c*_ = 0.3 cyc (Fig. 4A) and low ITDs (blueish colors) the input trajectories in the MSO-LSO plane have zero angle relative to the MSO axis, whereas high ITDs (reddish colors) tend towards steeper positive angles. The difference in angles is more pronounced the higher BF. The angle in the MSO-LSO plane thus encodes stimulus ITD and can be read out by a (normalized) synaptic weight vector

**Fig 4.**
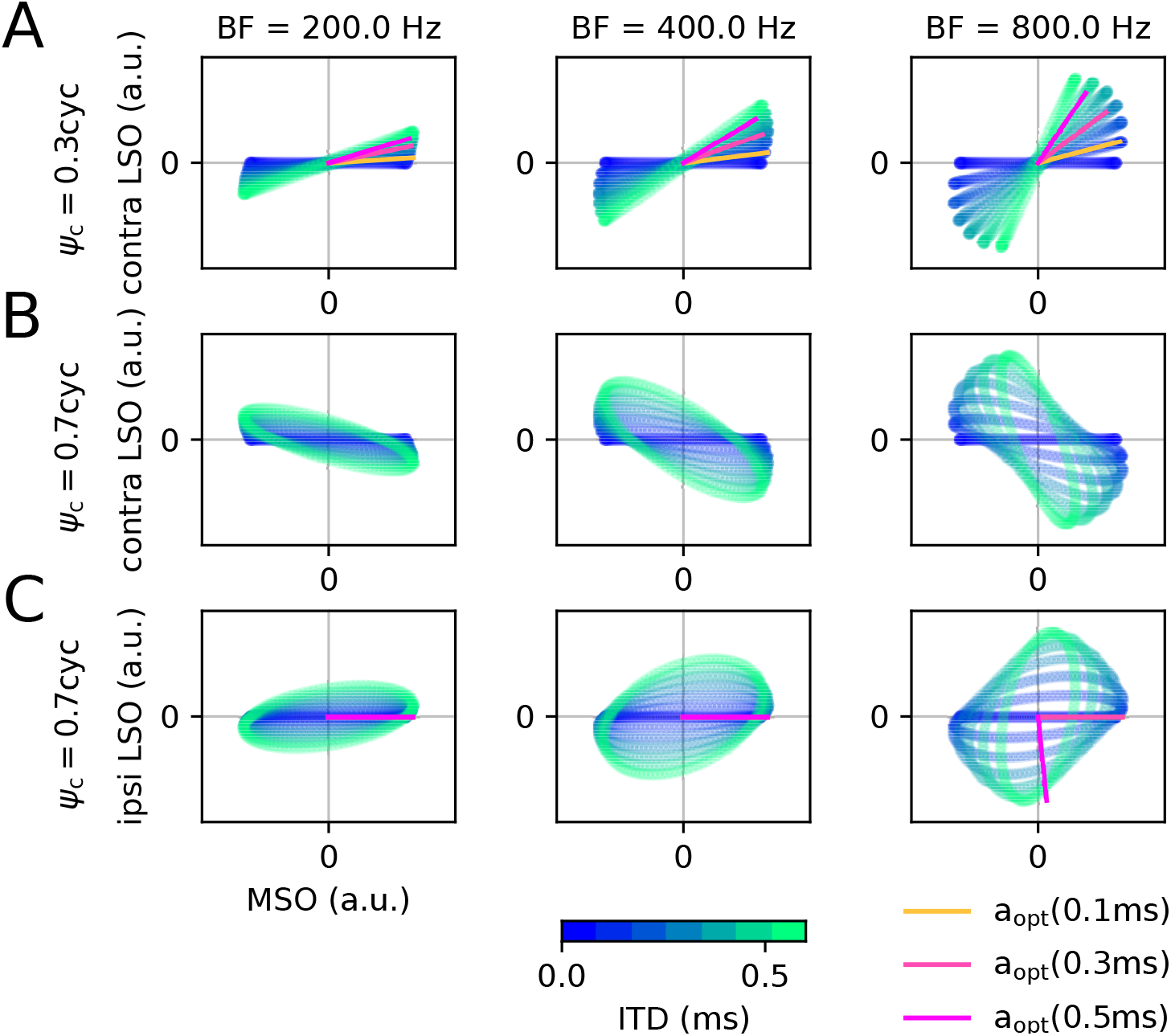
Brainstem code. The input space (LSO vs. MSO) is plotted for three BF channels (columns as indicated), when stimulated with a pure tone at BF. (A) For a commissural phase *ψ*_*c*_ = 0.3 cyc the ITD (blue to red colors) changes the inclination of the trajectories. The optimal weight *a*_opt_ from Fig. 3 reflects the direction in MSO-LSO space (greenish color) that matches the inclination of the corresponding ITD. (B) For *ψ*_*c*_ = 0.7 cyc ITDs are reflected by negative inclinations, which could only be sampled by negative weights *a*, but are forbidden since the contralateral LSO is excitatory. (C) Using the inhibitory ipsilateral LSO as a second input dimension again results in a positive inclination for *ψ*_*c*_ = 0.7 cyc, however, being an inhibitory projection, only negative weights *a* are allowed. The green directions reflect the optimal (non-positive) *a* maximizing the response amplitudes.

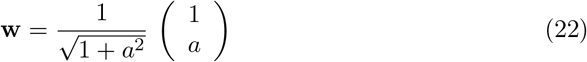

adjusted to a specific ITD (greenish lines in Fig. 4) using the optimal *a* parameters from Fig. 3. For non-optimal *ψ*_*c*_ = 0.7 cyc (Fig. 4B) the ITD separate towards negative angles in the input space. Since, however, the contralateral LSO is excitatory, negative relative weights *a* cannot be established. Also taking into account the ipsilateral LSO activity (Fig. 4C), does not solve the problem: the ITDs now separated along positive angle in the MSO-LSO space, however, the ipsilateral LSO exerts inhibitory derive to IC neurons and thus the relative weight *a* cannot be positive. The optimal weight is thus largely *a* = 0, except for very high ITDs and high frequencies, at which strong negative weight evokes the optimal IC response.

Thus, the optimal relative LSO weight *a* in each frequency channel allows an IC neuron to tune its maximal responsiveness to a specific ITD. The optimal commissural phase *ψ*_*c*_ determines the shape of the trajectories in MSO-LSO space. For *ψ*_*c*_ ≈0.3 cyc, the trajectories are confined to positive angles that increase with ITD. A further advantage of decoding via an angle in MSO-LSO space is level-invariance: Scaling both signals by the same factor (i.e., making the sound louder or softer) would not change the angle and thus also not require a different decoder.

So far the model is designed entirely feed forward. The IC, however, also hosts recurrent connections [26]. In this paper the recurrent IC network is assumed to provide inhibition within a BF channel such that the activity of all neurons is suppressed according to the total amount of activity in the BF channel. To this end, IC_*τ*_ (*ω*) is considered as the input to the IC neuron at target ITD *τ*, which yields the output activity 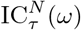 by normalization with the mean input

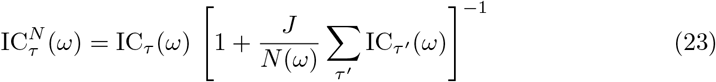

with *N* (*ω*) denoting the number of neurons with BF=*ω/*(2*π*). All simulations were performed for an inhibitory synaptic weight of *J* = 20. The main effect of inhibition is thus to sharpen the activity peak along the target ITD (*τ*) axis. In this paper, inhibition is only included to firing rate computations, but not to measures based on sound reconstructions (Pearson’s correlations), which require linearity.

For the final set of simulations, we also included rate adaptation, implemented by attenuation variables *α*_MSO*/*LSO_ *<* 0 that multiply the corresponding IC inputs *MSO*(*t*) and *LSO*(*t*) from eqs. 8. The attenuation variables follow first order kinetics with steady state 1, i.e.,

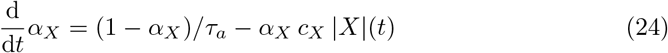

where *X* ∈ {MSO,LSO}, and *X*(*t*) refers to *MSO*(*t*) or *LSO*(*t*), respectively. The adapted SOC activities are then obtained by replacing *X*(*t*) by

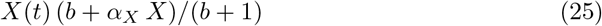

in which *b/*(*b* + 1) implements a minimal baseline activity level for the maximally adapted state *α*_*X*_ = 0. Simulations have been obtained for *b* = 0.1, *τ*_*α*_ = 2s, *c*_*MSO*_ = 5*/τ*_*α*_, and *c*_*LSO*_ = 10*/τ*_*α*_ (with the input signal normalized to maximum amplitude 1).

### Simulation details

The IC model outputs were computed in Fourier space with frequency dependent parameters *a* and *ψ*_*c*_ from Fig. 3. In detail, left and right ear sounds were sampled with 96 kHz. Broad band sound stimuli were low-pass filtered by a first order butterworth filter with cutoff frequency 500Hz unless otherwise mentioned. This filtering reflects processing in the low-frequency pathway, and particularly the limits imposed by the synaptic integration time constant at IC neurons of about 2 ms [26]. A fast Fourier transform was applied to snippets of 4001 samples enveloped with a Blackman window. Subsequent snippets were offset by 2000 samples, such that the overlap of snippets was approximately 50%. The fast Fourier transformed sounds for the left and right ear were then mapped to an IC response according to Eq. (15) in each frequency channel. After back transformation into the time domain the windowed snippets were added up with appropriate time shifts corresponding to the 2000 sample points.

The delay-line model was simulated according to eq. (5).

### Stimuli

All sound stimuli used in this study were sampled at 96 kHz. We used a) pure tone stimuli with frequencies of 200, 300, 400, 600 and 800 Hz, b) broad band white noise, c) speech signals of two male and one female speaker reading passages from free audiobook. The speech excerpts are available as Supporting Information S1. For tone in noise detection experiments, sound levels were adjusted such that 0dB corresponds to the same root mean square (RMS) of the 400Hz pure tone and a noise filtered with a 2nd order butterworth band-pass filter with width 200Hz centered around 400Hz.

## Results

First, single cell ITD tuning curves and best ITDs were checked for consistency with experimental data. Although the model was not optimized for generating a broad coverage of ITDs, but rather to maximize firing rate for any given ITD (which still would allow an even higher rate for a different ITD), tuning curve peaks cover the full physiological range (here 0.3ms) and beyond (Fig. 5A). Best IPDs are almost constant over BF (Fig. 5B) with mean best IPD at around 0.125 cycles (Fig. 5C) as found experimentally [17]. Hence, although IC neurons integrate LSO and MSO input with additional commissural delay, the distribution of best IPDs matches the empirically found contralateral bias. Expectedly, the shapes of the ITD tuning curves (particularly for lower BFs) deviates from physiological data (e.g., [64]), since the model output |*IC*| ^2^ rather corresponds to membrane potential amplitudes and the spike rates would be a non-linear transformation of |*IC*| ^2^. The Best ITD, however, would be rather unaffected by such a monotonous non-linearity.

**Fig 5.**
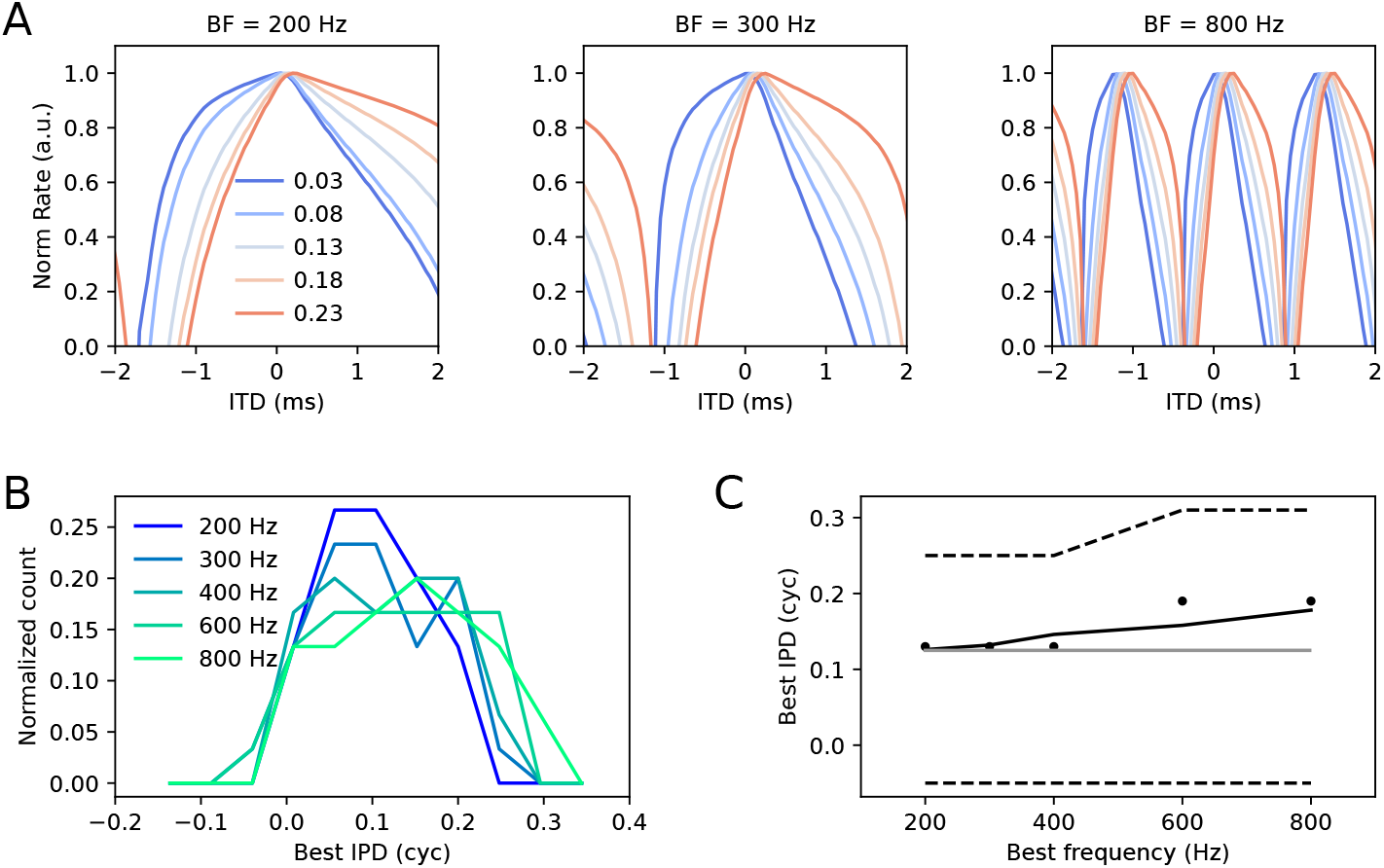
Best IPDs. (A) Example tuning curves in the 200, 300 and 800 Hz channels obtained with pure tone stimulation at BF. Different colors indicate responses of neurons with different Target ITDs (in ms). Outcomes from equation (19) are rectified at a threshold of 0.67 times the population mean in the physiological range to mimic a non-linear spike threshold. (B) Distribution (normalized to sum 1) Best IPDs for BF channels as indicated by color. Best IPDs were derived from the phase of the Fourier mode at BF. (C) Mean (solid), median (dots), and 10 and 90-percentiles (dashed) of best IPDs. Grey line indicates 0.125 cycles. Tuning curves were obtained for a physiological ITD range of 0.3 ms; middle panel in Fig. 3.

Next, the model was tested on its capability to follow dynamically varying ITDs [39], although the parameters were not optimized for ITD encoding. For comparison with experimental data from [24], “phase warp” stimuli were applied, which are a broad band version of a binaural beat that evoke a strong impression of acoustic motion for low warp frequencies *f*_warp_ and binaural flutter for *f*_warp_ ⪆ 5 Hz. The stimuli were created in frequency space such that for one ear, each frequency (*f*) component was assigned equal magnitude and a random phase *ϕ*_*f*_, and for the other ear that phase *ϕ*_*f*_ was assigned to the frequency component shifted upward by a difference *f*_warp_. The simulated IC population activity indeed reflected the fastly fluctuating ITDs (Fig. 6A) and also the broad-band nature of the stimulus (Fig. 6B), as described psychophysically and physiologically [24, 38, 39]. Phase-locking to the warp frequency, as measured by vector strength,

**Fig 6.**
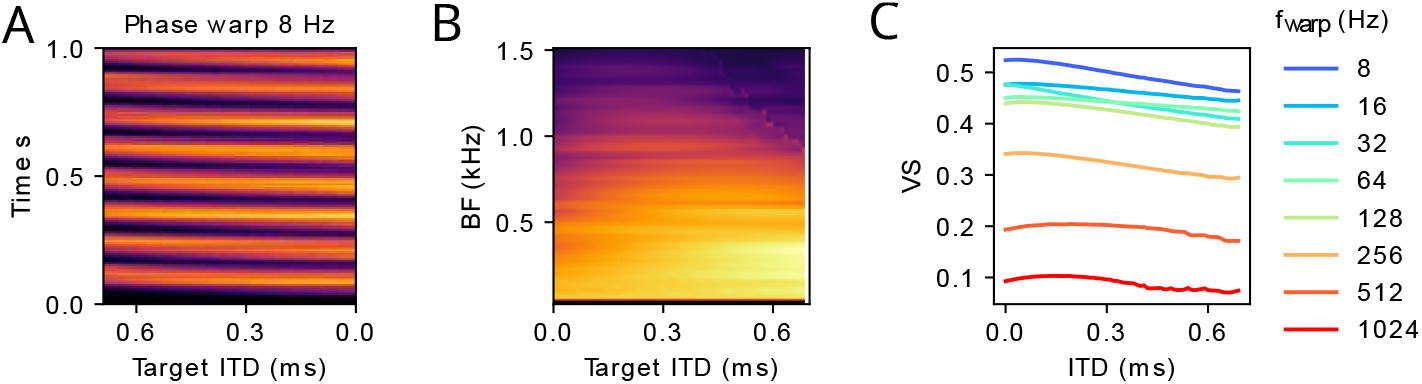
Fast ITD sensitivity. (A) Population rate as a function of time separated into neuron populations optimized towards the same ITD and averaged over all frequency bands. Color code was normalized between 0 (dark) and maximum (bright). (B) Same simulation as in A averaged over time instead of BF (same 0-max color code). (C) Phase-locking to warp frequency as a function of target ITD and warp frequency (color) measured by vector strength (VS).

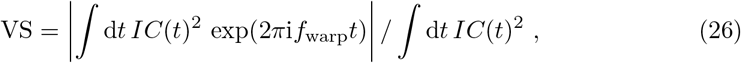

was observed up to 512 Hz, similarly as reported from electrophysiological measurements [24], reflecting the 500 Hz low-pass filtering that is supposed to model the postsynaptic current kinetics. Also the numerical VS values roughly matched experimental values of [24] in their Figure 5.

After supporting the model design by comparison with dynamic ITD processing, a classical binaural detection paradigm was applied: Binaural unmasking has been defined as the improvement of signal detection thresholds by presenting signal and masker with different interaural phases (as compared to presenting them with the same phase at both ears), without additional amplitude cues (as e.g., ILDs or gap listening) [37]. The difference in detection threshold is called binaural masking level difference (BMLD). To generate interaural phase differences, traditionally either signal (S_*π*_N_0_) or masker (S_0_N_*π*_) have been sign inverted. In Fig. 7, the model outputs are evaluated for the classical S_0_N_0_ vs. S_*π*_N_0_ situation. A reduction of the signal-to-noise ratio (SNR) in the input selectively reduces the detectability of the tone signal at 400 Hz. For S_0_N_0_ the relative signal to noise differences *d*^max^ derived from the model output are more sensitive to stimulus SNR than for the S_*π*_N_0_ condition, suggesting that interaural phase difference improves tone detection thresholds, as shown in psychoacoustics [36].

**Fig 7.**
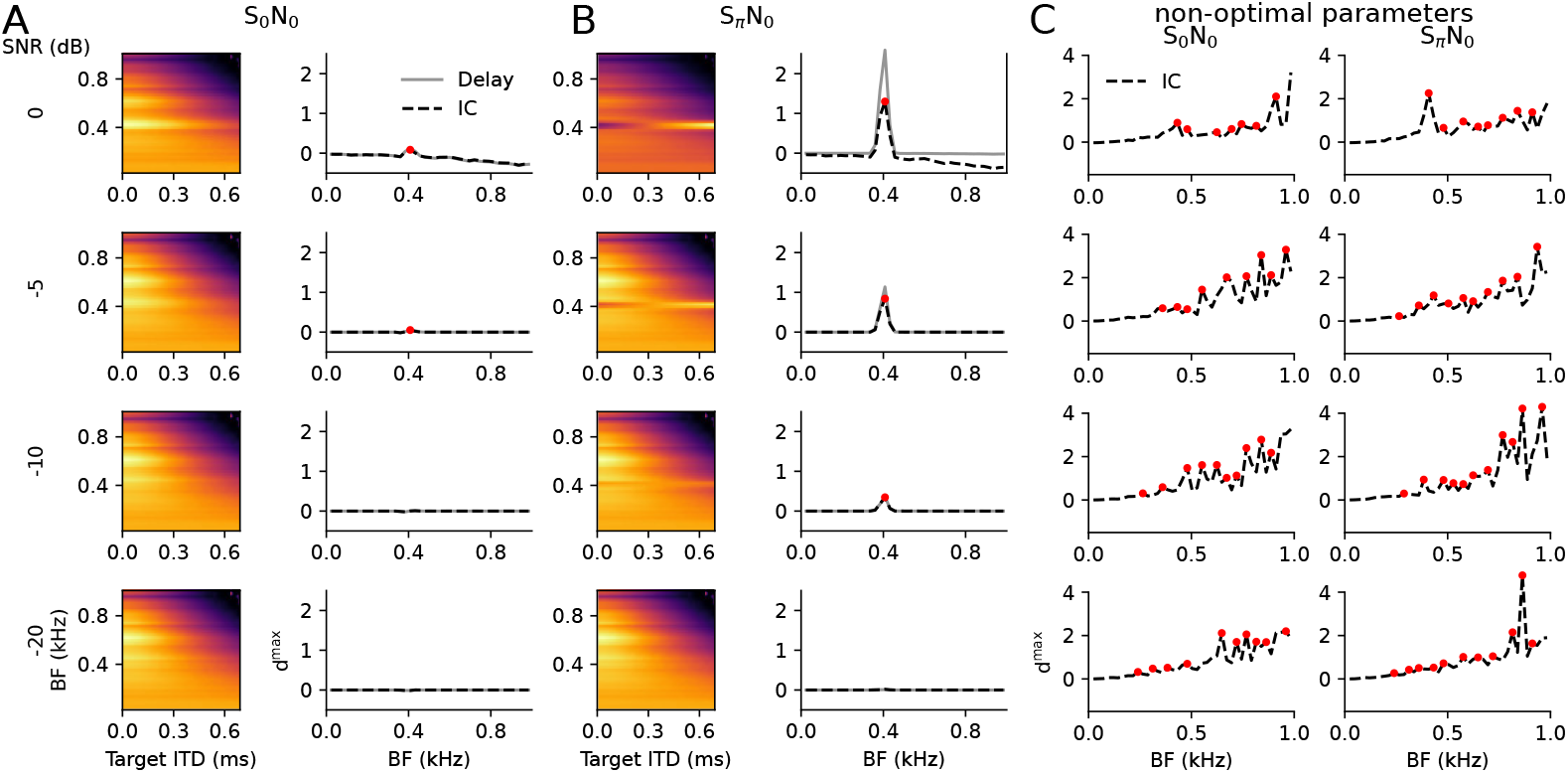
Binaural unmasking. A 400 Hz pure tone signal (S) and a white noise masker (N) are presented with binaural phase 0 for the noise and binaural phases 0 (A) and *π* (B) for the tone. The sound level of the signal is presented at different levels (rows), with SNR=0dB indicating that band-pass filtered tone and noise have the same intensity in a 200 Hz band centered at 400 Hz (see “Stimuli” paragraph in Methods). Left: Power |IC(*BF*) |^2^ of the IC model as function of BF channel and target ITD for which *a* was optimized to according to Fig. 3. The color code is normalized between 0 (dark) and maximum (bright). Right: Maximal normalized difference *d*^max^ between signal plus noise and noise alone in each BF channel. The results for the delay line model are depicted in grey. The red dot indicates the peak of IC model exceeding a prominence threshold of 0.05. (C) *d*^max^ plots for the two stimulus situations with a Gaussian jitter (*σ* = 0.2) applied to the optimal parameters *a* and *ψ*_*c*_.

While these simulations can provide a qualitative basis for the biological mechanisms underlying the BMLD, quantitative explanation of BMLDs are much harder, also because the perceptual thresholds involve additional higher-order processes that “read” the IC population pattern. BMLDs of about 15 to 20 dB are reported in psychoacoustic experiments [65], while BMLDs derived from single neurons are only few dB on average [66, 67]. Thus extracting psychoacoustic threshold requires additional statistical assumptions beyond the neural tuning properties. An estimate for detection thresholds that could be applied to our simulations is the peak prominence of the relative differences 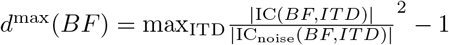 between model output from “tone plus noise” and model output “noise alone” as a function of BF. Peak prominence is defined as the peak amplitude minus the highest local minimum. Setting a prominence threshold of 0.05 approximately explains experimental BMLDs (red dots in Fig. 7) without attempting to fit all psychoacoustics data. For the delay line model (grey *d*^max^ curves) peak prominences in the S_*π*_N_0_ condition are generally higher, but the threshold of 0.05 would be crossed at comparable SNR values, yielding similar BMLDs. Notably, we are not proposing peak prominence as a psychophysical readout model, but rather a biologically plausible way of how a population code could be read by downstream neurons (potentially involving multiple stations).

In order to test the sensitivity of the BMLD effect on our optimization procedure (Fig. 3), we repeated the simulation with a 20% random Gaussian jitter around the optimal *a* and *ψ*_*c*_ values (Fig. 7C). Parameter jitter introduces spurious peaks of *d*^max^ and generally reduces detection thresholds beyond psychoacoustic values [65]. Detection peaks at high BF channels (with no signal power) indicate that a non-optimal choice of the parameters removes much of the binaural correlations and inhibition amplifies spurious correlations in the noise. Hence, deviations from the optimal parameters have marked effect on detection thresholds.

To furthermore evaluate the effect of inhibition, simulations were repeated in a pure feed forward setting (*J* = 0); Fig. 8. While response patterns between S_0_N_0_ and S_*π*_N_0_ conditions clearly separate in terms of target ITD (x-axis), the detection thresholds for low SNRs (between -25 and -30) seem much less distinct between S_0_N_0_ and S*π*N_0_ than for simulations with inhibition. Thus the model proposes BMLDs to fundamentally arise from negative feedback between neurons of distinct target ITDs.

**Fig 8.**
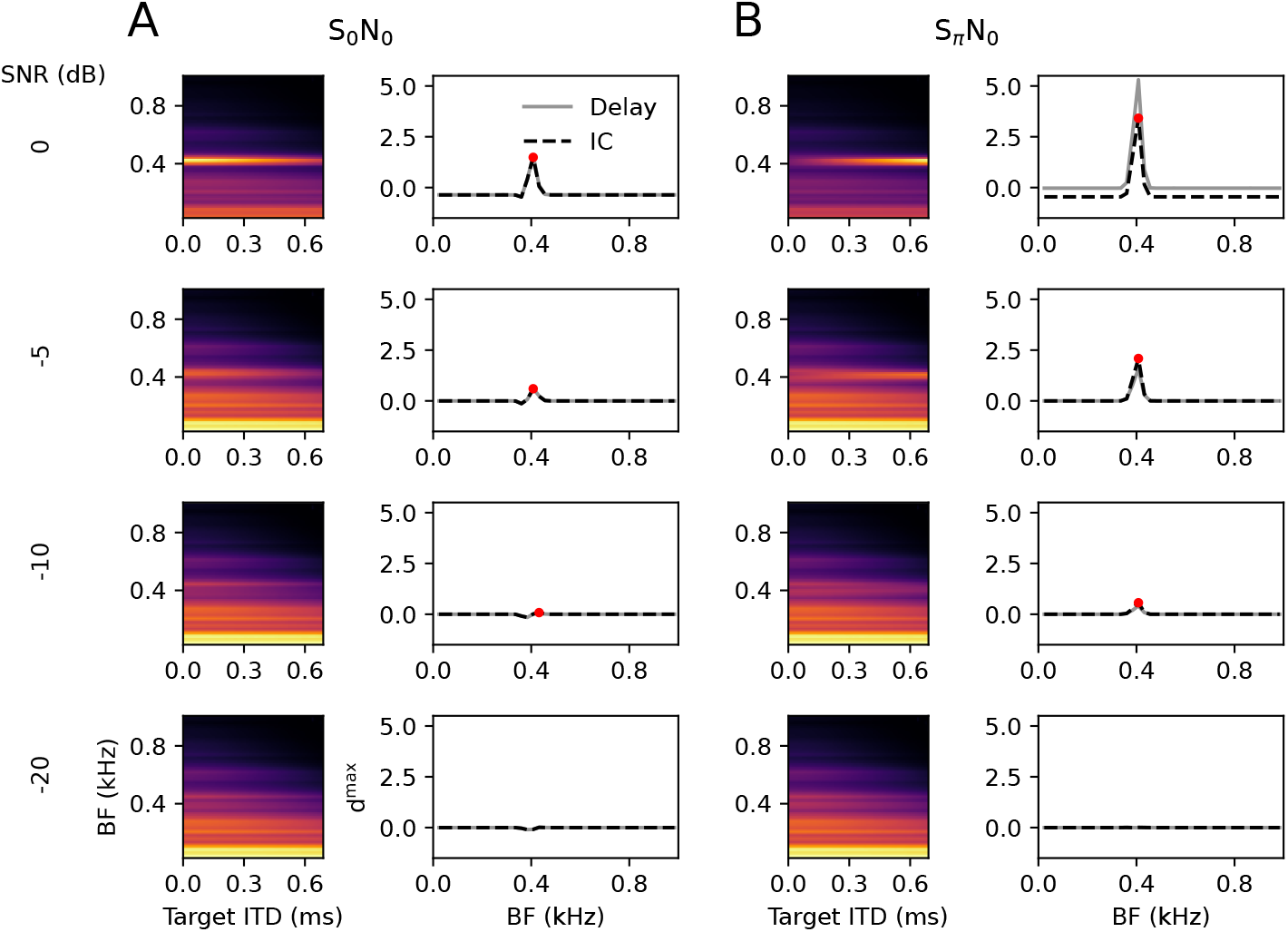
Binaural unmasking without inhibition. Same as Fig. 7 with inhibitory coupling set to *J* = 0.

To quantitatively summarize Figs. 7 and 8, we computed detection thresholds (relating to the red dots in the previous figure) as a function of prominence threshold and for various levels of inhibition (Fig. 9). Expectedly, detection threshold rise as a function of the prominence criterion. Psychophysical BMLDs of about 15dB (e.g. [65]) are better achieved for a high inhibition case (*J* = 40) (Fig. 9A), and also psychophysical detection thresholds for the S_*π*_N_0_ stimulation (−20dB and below) rather fit to the high inhibition case and very low prominence thresholds (Fig. 9A). Without inhibition (Fig. 9B), BMLDs are consistently below 10dB and thus unrealistic, further corroborating that intra-IC inhibitory circuitry may play an important role in sharpening of the peaks. To rule out that our model results arise from artificially removing peripheral filtering, we repeated the simulations with a variant of the model, in which the sounds are first filtered with a bank of 41 4-th-order gammatone [68] filters with equally spaced center frequencies between 24 and 984 Hz (500 Hz low pass is applied after peripheral filtering to account for synaptic filtering in the IC); Fig. 9C. Thresholds obtained with peripheral filtering are relatively consistent with results from Fig. 9A, except for high prominence thresholds where the inhibitory weight *J* seems less important. Again, psychophysically realistic thresholds require rather low prominence thresholds, suggesting that the IC population activity would be read out at high acuity.

**Fig 9.**
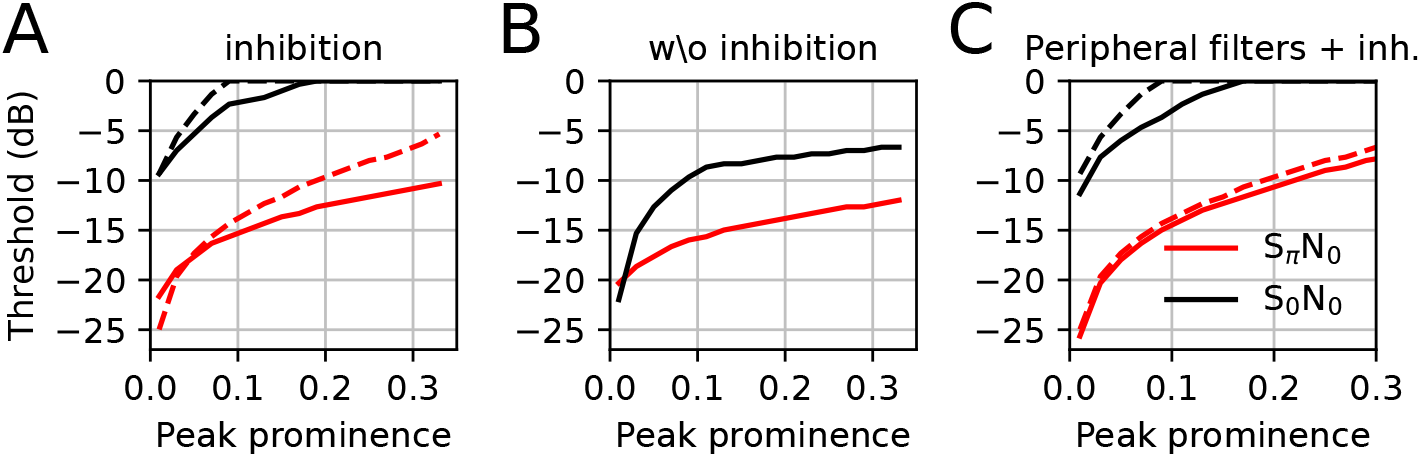
Detection threshold as a function of peak prominence parameter for S_0_N_0_ (black) and S_*π*_N_0_ (red) condition. Only stimuli with SNRs*<* 1 have been probed, thus no positive thresholds levels could be obtained. (A) Thresholds for model described in Fig. 7 (solid) and with double inhibitory strength *J* = 40 (dashed). (B) Thresholds without inhibition (Fig. 8, *J* = 0). (C) Thresholds derived from a model variant where sound stimuli were pre-processed by a gamma-tone filter bank (solid: *J* = 20; dashed: *J* = 40).

The classical S_0,*π*_N_0,*π*_ stimuli can be criticized as unphysiological, since an interaural phase difference of *π* does not correspond to a real sound location for frequencies below approximately 700 Hz. Also for wide band noise maskers, as well as broad band natural sounds, a sign inversion cannot be mapped to a position in space. Binaural unmasking, however, was also shown to exist for ITDs [69] representing a physiologically more realistic sound separation scenario. Therefore, in Fig. 10, sound sources were assigned a sound location via a physiological ITD, 0 and 0.4ms, as indicated in Fig. 10.

**Fig 10.**
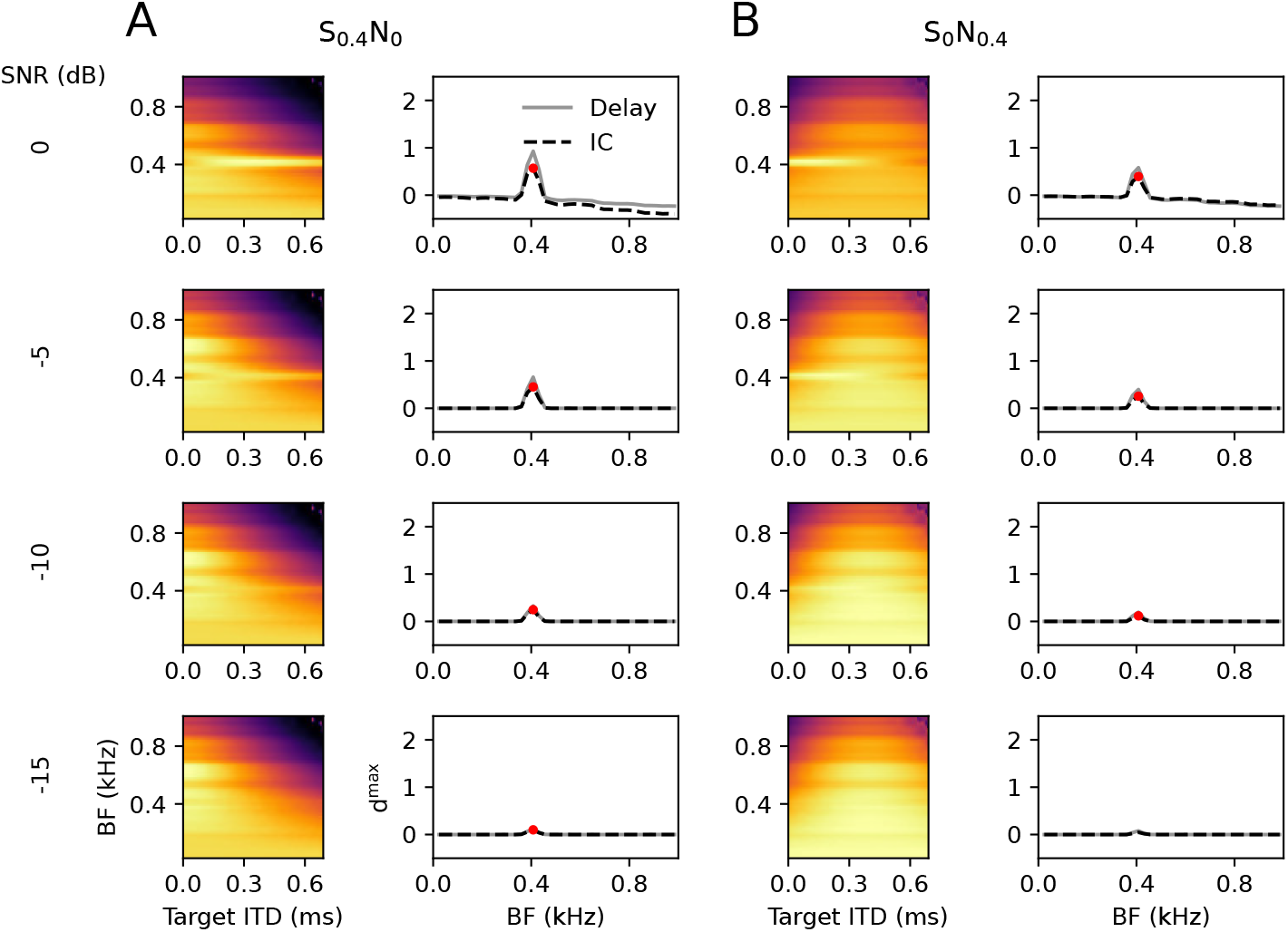
Binaural unmasking with physiological ITDs. Same as in Fig. 7. Note the SNR scale is reduced as compared to Figs. 7 and 8.

Consistent with the literature [7] the threshold for S_0_N_0.4_ is higher than that for S_0.4_N_0_.

All results so far have been derived for low-pass filtered stimuli in order to reflect auditory processing in the low-frequency brainstem pathway. The theoretical derivations, however, at no point require a restriction to low frequencies. Particularly the delay model implements optimal reconstruction independent of the spectral content of the signal. To show that the IC model does not critically rely on the specific cutoff frequency of 500 Hz, stimulus separation, as measured by Pearson correlation model between output and signal source, was next tested for mixtures of 2 and 3 human speakers without low-pass filtering (Fig. 11). Correlation coefficients between individual speech sounds and model outputs in 20 ms windows revealed high reconstruction acuity in temporally alternating windows relating to one of the speakers (Fig. 11A). Both the IC model and the delay line model exhibited similar pattern, suggesting that the auditory midbrain circuitry achieves robust almost optimal sound separation performance independent of the specific phase-locking limit and the number of noise sources. The firing rate pattern across the population of neurons with varying target ITDs reflects the spatial locations in the contralateral hemisphere (Fig. 11B). Sounds with ITDs corresponding to the ipsilateral hemisphere evoke largest rates at target ITD 0. For acoustic verification, audio samples of the reconstructed wave forms, the mixtures, as well as the original sounds can be found as Supporting Information S1. At this point, however, it is important to note that fine structure ITD sensitivity (particularly in the LSO) is generally not observed above 1 to 2 kHz. The generality of the model results demonstrated in Fig. 11 suggests that high-frequency binaural processing could be performed in the same circuit if it was extended to envelope ITD sensitivity. This speculation, however, still requires more specific physiological and modeling support.

**Fig 11.**
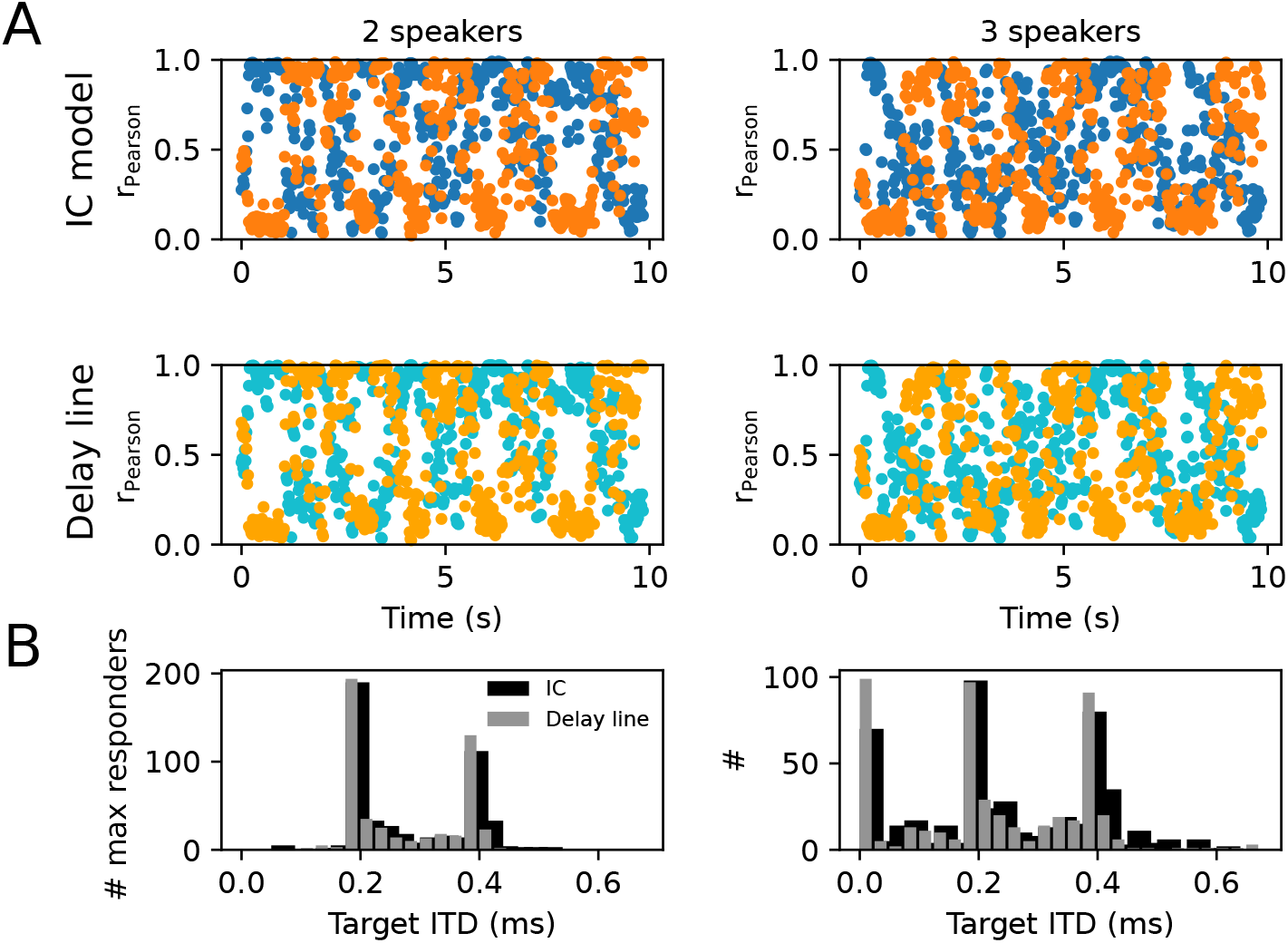
Deciphering a speech scene. Sound scenes were constructed from two (left column: female ITD 0.2ms, male ITD 0.4ms) and three (right column: same as left plus one additional male at ITD -0.2ms) speech signals at physiological ITDs. (A) Pearson’s r values were computed between the first two isolated speech sounds (female: blue, male: orange) and the broad band reconstructed IC model output [top row according to Fourier back transform of eq. (11)] and the delay line model [bottom row according to eq. (5)] in snippets of 20ms. Dots represent the mean Pearson’s r averaged over all target ITDs (0 to 0.7ms). (B) Histograms of how often a neuron with a given target ITD had the maximal response among its peers (with other target ITD) in a 20ms snippet (IC model: black; Delay line model: grey). Speech signals were taken from a free audiobook.

In a final set of simulations, the model was extended by an adaptive mechanism (see Methods), assuming that the SOC inputs show firing rate adaptation as reported from measurements [70, 71]. Simulation outcomes are shown in Fig. 12 for a pure tone stimulus and a female speaker applied to two contralaterally leading ITDs. Owing to the adaptation mechanism, over time, the angle of the input trajectories in the MSO-LSO plane moves towards the diagonal (Fig. 12A), which generates a trend towards balanced LSO and MSO activity (45^°^ angle in the MSO-LSO plane). In sounds with strong amplitude modulations such as speech (Fig. 12C) adaptation effects, however, decay during pauses and are no longer present at transients, indicating that ITD information is best preserved at onsets of transients, as reported psychoacoustically [72]. Such adaptation-induced fluctuations in neural activation recruits more neurons (i.e., neurons within a broader range of target ITDs) and thus could convey sound information with higher population rate during periods with little amplitude modulations at the cost of losing ITD information.

**Fig 12.**
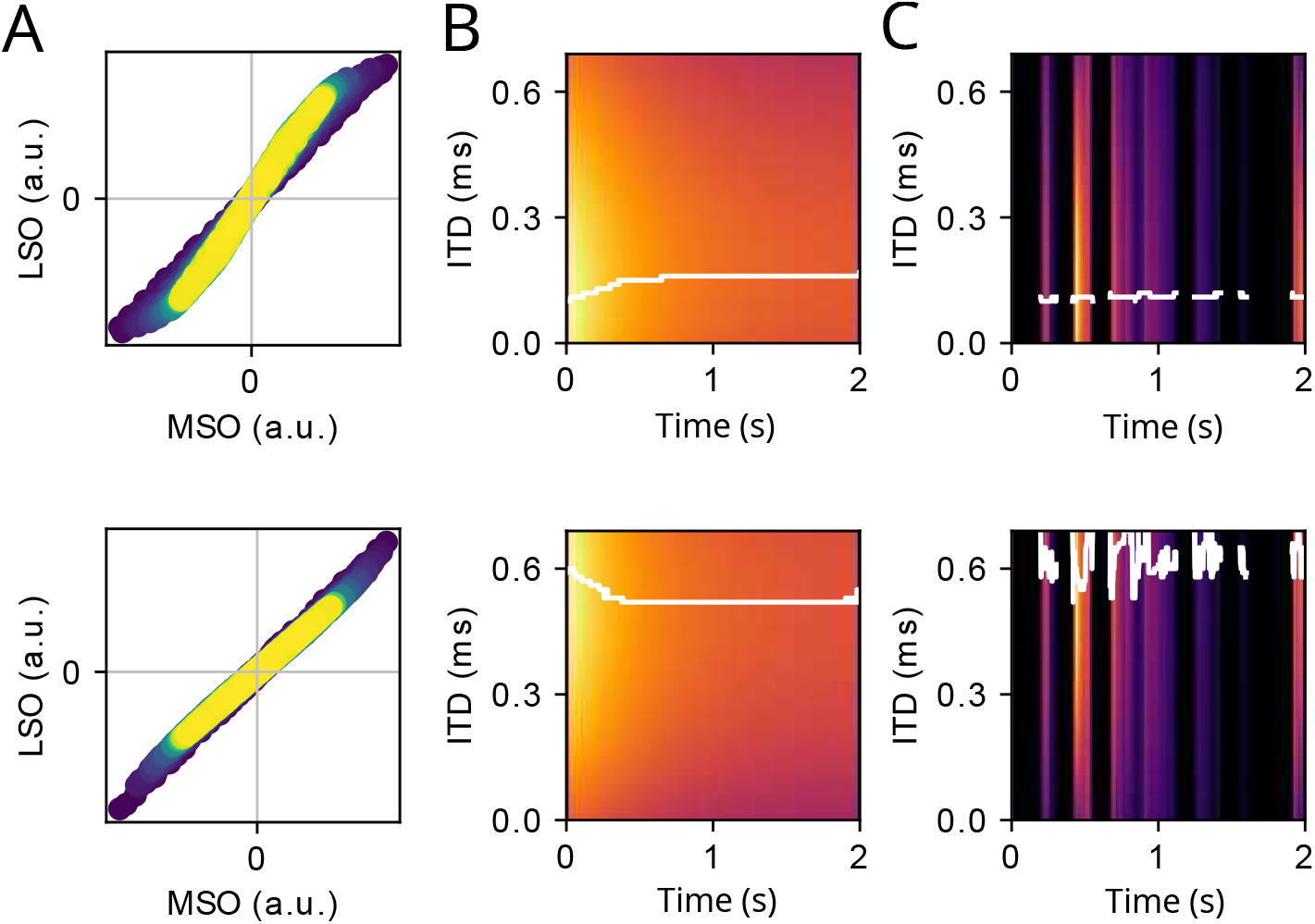
(A) MSO-LSO input plane (see Fig. 4) upon stimulation with a 2 second 400 Hz pure tone with ITD=0.1 ms (top) and ITD=0.6 ms (bottom). The color code reflects the stimulus time (blue early, yellow late). (B) IC population activity (target ITD of IC neuron on y-axis) stimulus ITD=0.1 ms (top) and stimulus ITD=0.6 ms (bottom). The color code is normalized between minimum (dark) and maximum (bright). The white transparent line marks the neuron with the maximum activity for any point in time. (C) Same as B for a speech sound from a female speaker.

## Discussion

A circuit-level model was presented on how the inferior colliculus processes binaural information computed in the superior olivary complex (SOC). For an optimal commissural phase delay *ψ*_*c*_ in the stria of Held of about 0.3 cycles, IC neurons can generate a broad range of best ITDs of the contralateral hemisphere by only adjusting the relative weight of the LSO input. Commissural phases far off 0.3 cycles cannot generate an ITD rate code in the IC given the physiological synaptic kinetics. The present model performs close to optimal in sound source reconstruction, as it closely matches the theoretically optimal performance of the delay-line solution. Generating diverse best ITDs via modulations of synaptic weights is biologically more efficient than using a range of delay lines, because it allows for dynamic adaptations on short time scales. Moreover, systematically varying delay lines have been shown to be absent in the mammalian brainstem [48].

The mammalian ascending auditory pathway leading up to the ITD sensitive IC neurons involves additional connections and nuclei beyond the basic circuit that has inspired our model (Fig. 1). Furthermore, there is species variability [73]. Most importantly, our model omits the dorsal nucleus of the lateral lemniscus (DNLL) [74], which also holds ITD sensitive neurons [75] and projects mostly GABA-ergic inhibition to the IC as well as to its contralateral counterpart. The DNLL has been implicated in echo suppression [76], but direct influence on ITD processing in IC cannot be excluded. We nevertheless refrained from adding these additional pathways to our model to first understand the implications of the most prominent projections in ITD sensitive mammals, and leave it for future work to understand potential functional consequences of the slower GABAergic DNLL inputs.

Moreover, ITD sensitivity of LSO neurons to low frequency tones has only been investigated little. The current best account [55], however, suggests high acuity of ITD encoding similar to MSO neurons. More data of low-frequency ITD sensitive LSO neurons is needed to support our model.

Current neurobiological circuit models of binaural hearing have a strong focus on explaining sound localization, including its mechanisms [62, 77–82], encoding [15, 30, 34, 35, 77, 83] and perception [31, 59]. Sound localization and binaural unmasking, however, involve the same neuronal brainstem structures MSO and LSO. Yet, biologically inspired models of binaural IC are scarce. As a notable exception, the work of Dietz et al. (2008) [39] has a strong focus on biological plausibility of the peripheral and SOC processing, however, makes no specific attempt to connect to the IC circuitry. The present model fills this gap in that it largely disregards the biological stages up to the SOC, but emphasizes the contribution of the feed-forward pathways to the IC. Our modeling approach also imposes limitations: 1) The effects of peripheral processing are reduced to Fourier decomposition into BF channels and a low-pass filtering of the signals to account for the phase-locking limit. Any non-linear effects of peripheral processes are thus neglected, specifically compression and half-wave rectification. Compression, however, could be compensated by level adaptation in the auditory nerve [84] and the brainstem [85] reducing the level dependence of firing rates. Half-wave rectification may be compensated for by the convergence of a large number of synaptic fibers at the level of the SOC neurons [56]. 2) Detection and localization thresholds cannot be quantitatively predicted by the model. Apart from the errors introduced by the missing peripheral model this is also because of the missing readout model, i.e., how the cortex translates the IC rate code into a percept.

The simplicity of the present model, however, is also its strength, as it has very little unconstrained parameters and as it is derived from the optimization principle of maximizing the response of an IC cell to a sound at a specific ITD. Particularly, avoiding a non-linear periphery model allows for a direct comparison of the reconstructed sound signal (without inhibition) with the original sound wave in the time domain and thus also a direct comparison with the delay line model, which also is formulated in the time domain.

Our model proposes two fitting parameters, *a* and *ψ*_*c*_, to be optimized to maximize the firing rate for a single sound stimulus at a given target ITD (under additional constraints that *a* is not too large, not always zero and predominantly positive). Optimizing *a* to maximize firing rate in an inhibitorily connected network is conceivable via Hebbian mechanisms in the feed-forward synaptic pathways implementing a winner-take-all type of receptive fields as suggested for visual maps [86]. Optimization of *ψ*_*c*_ does not need to occur on a single cell basis (because the optimal value 0.3 cyc is universal) and therefore could arise from general activity dependent myelin plasticity [87, 88]. The general circuit design of the mammalian auditory pathway, however, would also suggest that part of this optimization could have been achieved through evolution [73].

The present model furthermore resolves a shortcoming of two/four-channel models [27–29, 31, 33, 59], which so far do not provide a conceivable explanation for attentional effects. Mammals are able to focus their attention on a specific sound location of behavioral importance [89–91], however, neural mechanisms of attention typically suggest a bias towards neurons with receptive fields in the attended area [92]. Such a bias can be easily mechanistically implemented by facilitating the response or transmission of neurons with “attended” receptive fields [93], whereas it is hard to conceive such a bias for a push-pull code of space made out of relative spike counts in the whole neural population. Our model assumptions thus argue for a labeled line code arising in the IC due to synaptic mechanisms that then can be subject to attention-guided readout. Although the model would work across the whole broad-band spectrum, it is meant to apply only to low-frequency brainstem processing, since only there, phase-locked action potentials convey fine structure information of the sound wave form [94, 95]. Interaural level differences (ILDs) are not relevant for binaural unmasking at low stimulus frequencies but contribute sound separation and localization at high frequencies, particularly when ILDs are generalized to broadband short-term envelope fluctuations [37, for review]. While ILDs of high-frequency sounds are used for spatial hearing by all mammals, fine structure ITD information, is virtually absent in auditory nerve activity in mammals with exclusive high-frequency hearing or with very small head sizes. During evolution, however, early air-borne sound hearing mammals were such high-frequency hearers [96]. Also some contemporary bat species have too small heads to use ITDs, but nevertheless possess a monaural MSO [97]. Therefore the general assumption that the evolution of the MSO was driven by the demand to localize sounds seems not fully convincing and allows for the alternate hypothesis that the mammalian ITD circuit has evolved to perform (low-frequency)-sound source separation, with ITD-based sound location encoding evolving as a by-product. An extension of the present model to high frequency hearing (without fine-structure information) is a necessary next step to support this hypothesis.

## Supporting information

Supprting Information S1a

Supprting Information S1

## Supporting information

**S1 file. Sound files**. Zip files contains three original sound snippets female1.wav, male2.wav, male3.wav mixtures of two and three sources mixture.wav, mixture3.wav and reconstructions according to the delay line model (Jeffress) and the IC model. The original sounds were taken from a free audiobook (https://librivox.org/the-adventures-of-huckleberry-finn-version-5-dramatic-reading-by-mark-twain/)

## Acknowledgments

This work was funded by the Deutsche Forschungsgemeinschaft (DFG) under grant SFB 870 (TP B01). The authors would like to thank Benedikt Grothe for discussions.

## Code Availability

All Python code used to produce the results and the figures presented in this paper can be found under www.github.com/cleibold/ICmodel.

## Author Contributions

CL: Conceptualization, Formal Analysis, Funding Acquisition, Investigation, Methodology, Software, Supervision, Visualization, Writing – Original Draft Preparation; SG: Validation, Investigation, Methodology, Writing – Review & Editing

